# Age, wound size and position of injury – dependent vascular regeneration assay in growing leaves

**DOI:** 10.1101/2020.10.21.348680

**Authors:** Dhanya Radhakrishnan, Anju Pallipurath Shanmukhan, Abdul Kareem, Mabel Maria Mathew, Vijina Varaparambathu, Mohammed Aiyaz, Raji Krishna Radha, Krishnaprashanth Ramesh Mekala, Anil Shaji, Kalika Prasad

## Abstract

**Background:** Recurring damage to aerial organs of plants necessitates their prompt repair, particularly their vasculature. While vascular regeneration assay in aerial plant parts such as stem and inflorescence stalk are well established, those on leaf vasculature remained unexplored. Recently we established a new vascular regeneration assay in growing leaf and discovered the underlying molecular mechanism.

**Results:** Here we describe the detailed stepwise method of incision and the regeneration assay used for studying the leaf vascular regeneration. By using a combination of micro-surgical perturbations, brightfield microscopy and other experimental approaches, our new findings show that the regeneration efficiency decreases with aging of the leaf, and increases with the nearness of the wound towards the proximal end of the leaf.

**Conclusion:** This easy-to-master vascular regeneration assay is an efficient and rapid method to study the mechanism of vascular regeneration in growing leaves. It can be readily adapted for other plant species and can be combined with cellular and molecular biology techniques.

## INTRODUCTION

Due to their sessile nature, plants are frequently subjected to injuries caused by biotic and abiotic factors. These injuries when left unattended can compromise the plant immunity, growth and even survival (Hwang, Yu, and Lai 2017; Radhakrishnan et al. 2020). In order to overcome the adversities of wounding, plants evolved a remarkable repertoire of regenerative responses ranging from, wound healing in the form of local cell proliferation to complete replacement of amputated organs, such as root tip regeneration (Ikeuchi et al. 2016; Shanmukhan et al. 2020). Although numerous studies have probed the mechanisms underlying several regenerative responses in plants, investigation regarding regeneration potential of aerial organs are limited (Durgaprasad et al. 2019; Iwase et al. 2011; Kareem et al. 2015). Thus, despite their higher susceptibility to injuries than underground organs, there is a dearth of information on regeneration in aerial organs of plants, particularly, in the leaves. Although leaves play a crucial role in plant physiology, their regeneration potential has hardly been investigated (Kuchen et al. 2012; Radhakrishnan et al. 2020).

Leaves possess an elaborate network of vascular tissue with a central midvein that transports substances to-and-fro between the main plant body. Damages to the midvein calls for prompt repair, failing which the transport of substances, and consequently the growth of the leaf and its adjacent branch are impaired (Radhakrishnan et al. 2020; Sachs and Hassidim 1996). Recently, a new vascular incision assay in leaf was developed to study the wound repair and tissue restoration in response to injury. The assay revealed that the mechanically disconnected parental stands are reunited by regenerating vascular tissue that bypasses the site of injury. The assay was instrumental in understanding the molecular mechanism underlying vascular regeneration in aerial organs growing in normal developmental context. Upon injury a coherent feed-forward loop comprising of cell fate determinants, PLETHORA (PLT) and CUP-SHAPED COTYLEDON2 (CUC2) activate the local auxin biosynthesis leading to vascular regeneration in growing aerial organs (Radhakrishnan et al. 2020). Here we show that, in addition to the extent of the injury, regenerative ability of the leaf vasculature is determined by age of the leaf explant, and position of the injury along the proximo-distal axis of the leaf blade.

This easy-to-master, reproducible assay can be performed using readily available laboratory supplies. The convenience of performing real time confocal imaging and other molecular techniques such as quantitative real time PCR using the injured leaves makes the assay valuable in studying the molecular players and mechanisms regulating wound induced response and regeneration in the normal developmental context. The method will also be useful in studying the interplay between mechanisms of vein patterning during development and that of vein regeneration.

## MATERIALS

### Equipment

#### Equipment for in vitro culture

- Laminar Air Flow chamber (LAF)
- Sterile pipette-tips (200 µl and 1 ml)
- Micro pipettes
- 1.5 ml micro-centrifuge tubes
- Sterile disposable square Petri plates, size: 120 mm × 120 mm (Himedia PW050-1)
- Clingfilm (Himedia Phytawrap)
- Plant growth chamber (Percival AR-100L3).

#### Equipment for incision and sample collection

- Fine pointed tweezer (Dumont tweezer, Style 5)
- Sterile razor blade
- Forceps
- Microscissors-Vannas scissor straight (Ted Pella, 1340)
- Gloves
- Face mask
- 70% ethanol
- 35mm round petriplate

#### Equipment for microscopy

- Steriozoom microscope (Zeissstemi 2000) for incision and sample collection
- Confocal laser scanning microscope (Leica TCS SP5 II) for brightfield imaging
- Microscope slides (Labtech)
- Microscope cover glass 22×22mm (Corning 2850-22)
- Watercolour brush (with small bristles)

### Reagents for Seed sterilization

- Seeds of wildtype (Columbia) *Arabidopsis thaliana*
- 20% sodium hypochlorite
- 70% ethanol
- Autoclaved Milli-Q water.

### Murashige and Skoog (MS) medium preparation

To prepare 1L (Half-strength) MS medium, add 2.165 g MS salt (Sigma Aldrich, M5524) and 10 g sucrose (Sigma Aldrich, S0389) to about 850ml Milli-Q water. Adjust pH to 5.7 with 1 N KOH and make up the volume to 1L. Add 8 g plant Agar (Sigma Aldrich, A7921). Autoclave the medium (121 °C for 20 min) and cool it to about 45–50 °C. Add 1 ml of 100 mg/ ml filter-sterilized Ampicillin (final concentration in medium-100 µg/ml) to 1L medium and pour 50 ml into each sterile square Petriplate (Himedia, PW050) within the LAF. Allow to cool and solidify.

### Reagents for decolourising sample

- 15% ethanol
- 50% ethanol
- 70% ethanol
- 96% ethanol
- 100% ethanol
- Glycerol (Sigma Aldrich, G5516)
- Chloral hydrate (Sigma Aldrich, 23100)
- Milli-Q water

Preparation of clearing solution: Dissolve 8g chloral hydrate in 3ml water. Vortex the solution until chloral hydrate is completely dissolved. Add 1ml glycerol to the solution.

Note: The solution has to be freshly prepared for clearing the leaf samples. It is also used for mounting samples for brightfield confocal microscopy.

## METHOD

### Seed sterilization

Seed sterilization should to be performed within LAF under sterile conditions. The work space and tools (Micro pipette, tip boxes, reagent bottles) required for the procedure have to be thoroughly wiped with 70% ethanol and UV irradiated. Prior to commencement of *in vitro* culture, hands should be washed using soap and wiped with 70% ethanol. A liquid surface sterilization protocol for seeds is described here.

1. Aliquot the required number of wildtype seeds in 1.5ml microcentrifuge tube. Note: For efficient sterilization do not take more than 300 seeds per tube and remove any debris such as parts of siliques left over from seed collection.
2. Add 1ml 70% ethanol. Agitate the contents by inverting the tube for 2-3 minutes. Note: Prior to centrifugation, ensure that the centrifuge and rotor surfaces are clean. Avoid touching inside the lid of the microcentrifuge tube while opening and closing to minimize contamination.
3. Brief spin the tube at 6000rpm and carefully discard the ethanol without losing any seeds.
4. Add 1ml 20% sodium hypochlorite and shake the contents for 2-3 minutes. Repeat step 3.
5. Wash the seeds 5-7 times using 1ml sterile autoclaved Milli-Q water.
6. Stratify the seed in 1ml sterile autoclaved Milli-Q water for two days at 4°C.
7. Plate the seeds on half strength MS medium in rows with at least 0.5cm gap between consecutive seeds. Note: For the ease of incision, avoid placing the seeds very close to each other.
8. Incubate the petriplates vertically in growth chamber under 45 μmol/m^2^/s continuous white light at 22°C and 70% relative humidity.

### Leaf Incision

1. Leaf incision can be performed on work bench after adopting necessary measures to minimize contamination. Wear gloves and face mask during the procedure. Prior to the incision, wipe the surface of the dissection microscope (Zeiss stemi 2000) and gloved hands with 70% ethanol. The tweezer for incision should be dipped in 70% ethanol and left to dry for a few minutes before incision. Opening the plate on multiple days for incision will increase the incidence of contamination. Note: Due to the fragile nature of the tweezer tips, sterilization techniques that may damage it or make it blunt are not recommended.
2. The plate containing seedlings is opened under a dissection microscope to confirm the age of the seedling and to perform incision in 5dpg (days post germination) old seedlings. Note: Due to the asynchronous nature of seed germination, all the seedlings may not be of the same age. To maintain consistency, only injure plants which are at the desirable developmental stage. Move aside the uninjured plants to distinguish them from injured ones. Age of the seedlings is important as older seedling display reduced regeneration efficiency while very young leaves are extensively damaged during the procedure. The appropriate age of incision is between 4-6dpg (Figure1).
3. Out of the two leaves belonging to the first pair (true leaves), the leaf that faces the lid of the petriplate is chosen for incision due to the ease of access. Using the sharp tip of the tweezer an incision is made on the lower abaxial surface of the leaf belonging to the first pair. The incision is carefully performed at the junction between petiole and basal end of the lamina (Fig. 1A). This ensures that the injury occurs just above the first lateral vein (counted from base of the leaf) where the regeneration efficiency is highest in comparison with other positions along the proximo-distal axis of the midvein (Fig. 2). The incision should be performed with just enough force so that it punctures the vascular tissue located close to the abaxial surface of the leaf without piercing out through the adaxial surface. This is important as extensive damages are not repaired (Radhakrishnan et al. 2020)(Fig. 1P). Note: Care should be taken, not to inflict multiple damages on the leaves. Therefore, it is not advisable to perform incision in both leaves of the first pair. During incision, a 200ul sterile microtip or sterile forceps can be used to restrict the movement of the plant. However, avoid touching the damaged leaves as it can inflict further damage.
4. After incision, the plates are closed and incubated vertically under continuous light condition at 22°C in growth chamber.
5. Four days post incision (dpi), the injured leaf is carefully cut at the petiole using a Vannas scissors and placed in 15 % ethanol in small round petriplates (35mm). Note: Around 20-30 leaves can be treated using 2-3ml ethanol in 35mm petriplates during the decolourising procedure. Alternatively, 6 well plates can be used when handling multiple samples.

**Figure 1:**
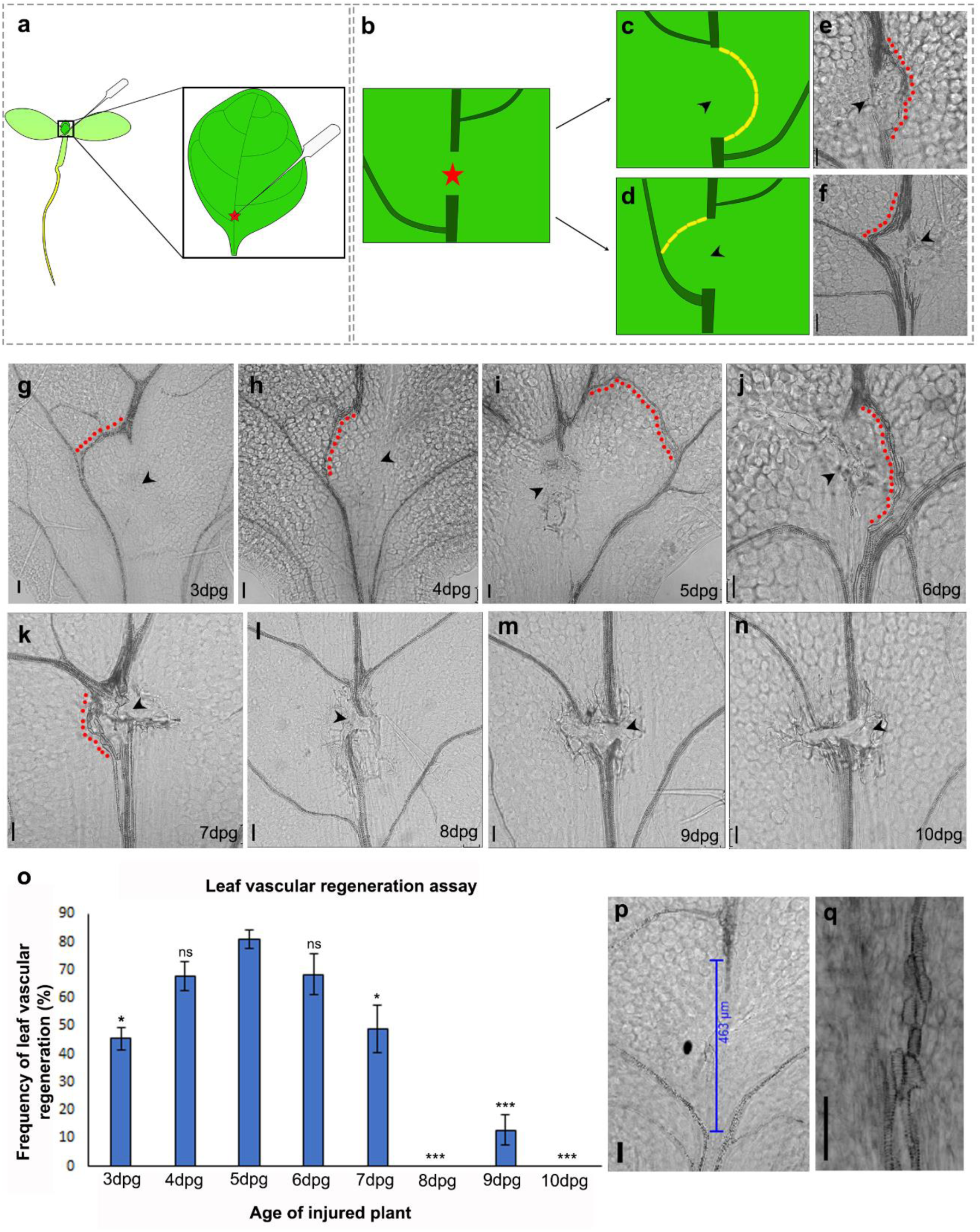
Leaf vascular regeneration upon midvein injury depends on the age of the injured leaf. (a-b) Illustration depicting the location of incision for effective vascular regeneration. Red star represents site of injury. (c,e) Schematic and brightfield image show the regenerating vasculature re-uniting the disconnected parental stands forming a D-loop bypassing the site of injury. (d,f) Illustration and brightfield image show the regenerating vasculature connecting to the nearest lateral vein. Yellow blocks in the (c) and (e) represent end-to-end connected xylem elements. (g-n) Regeneration response in leaves injured at 3dpg (g)(\**P*=0.016, n=24), 4dpg (h)(*P*=0.426,not significant (ns), n=45), 5dpg (i)(n=20), 6dpg (j)(*P*=0.605,not significant (ns), n=40), 7dpg (k)(\**P*=0.03, n=34),8dpg (l)(\*\*\**P*=1.6 × 10^−12^, n=43), 9dpg (m)(\*\*\**P*=2.405 × 10^− 9^, n=52), and 10dpg (n) (\*\*\**P*=4.8 × 10^−08^, n=21). Statistical analysis by Pearson’s χ^2^ test. Note that the 3-7dpg leaves are capable of reconnecting their disconnected vasculature but the regeneration efficiency declines with progressive aging of leaves. The regenerating vasculature is indicated by red dots. Black arrowheads indicate the site of injury. (o) Graph depicts frequency of leaf vascular regeneration in leaves injured at different ages. (p) Extensive damage creates a gap exceeding 400µm between parental vascular strands, as a result no vascular regeneration is observed. (q)End-to-end attached xylem elements of regenerated vascular strand. Scale bars represent: 50µm

**Figure 2:**
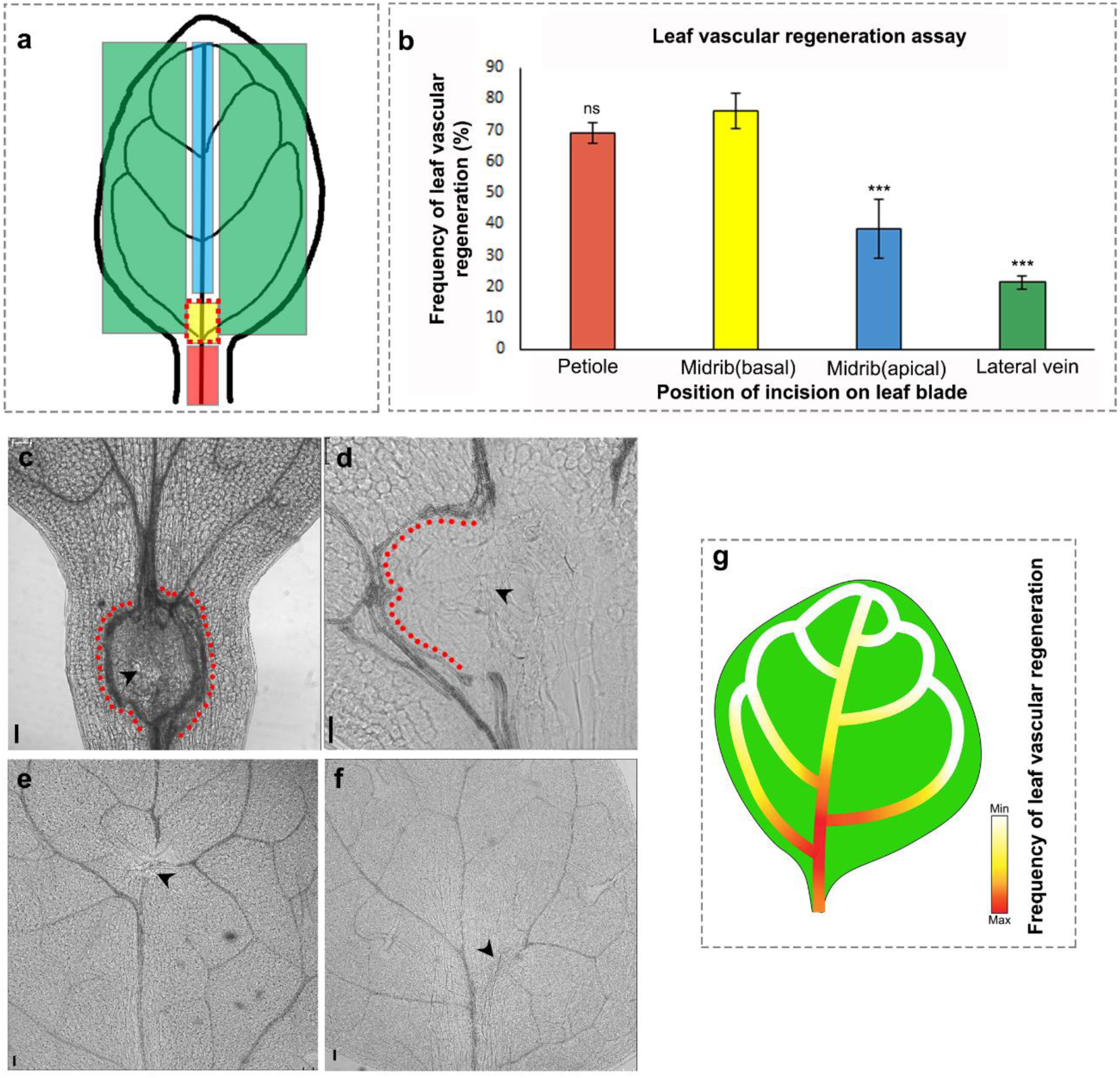
Vascular regeneration in leaf depends on the position of injury in the leaf. (a) Schematic depicting positions of incision on the leaf blade and petiole. (b) The frequency of vascular regeneration at different positions (represented by coloured boxes in (a)) of the leaf are represented by the same colour bars in the graph (b)Petiole (n=21, *P*=0.95, not significant [ns]), basal correct position (n=45), midvein upper end (n=42, *\*\*\*P*=0.0009), lateral vein (n=21, *\*\*\*P*=0.0002). (c-f) Images show the incision to vasculature in petiole (c), base of midvein (d), apical region of midvein (e), and lateral vein (f). Note the multiple strand formation upon injury in petiole (c). Red dots indicate regenerated vascular strand and black arrowheads represent site of incision. Scale bars represent:50µm. (g) Gradient represents the efficiency of vascular regeneration along the leaf blade with maximum regeneration (represented by red) at the base of mid vein. Lateral veins and distal end of midvein exhibit reduced regeneration frequency.

### Decolourising and clearing samples

1. The sample is placed in 2-3ml of 15% ethanol for 15 minutes. Note: Initially the leaves float on the surface of the solvent. Gently submerge the leaves using a paint brush with small bristles.
2. The ethanol is drained using micropipette with care being taken not to damage the samples.
3. Similarly, the samples are treated with 50%, 70% and 96% ethanol consecutively for 15 minutes each. After discarding 96% ethanol, the leaves are incubated for 12 hours in 100% ethanol for dehydrating the tissue and to remove chlorophyll pigmentation.
4. After discarding the ethanol, the samples are consecutively incubated for 15 minutes each in 96%, 70%,50% and finally 15% ethanol to rehydrate the samples.
5. After discarding the ethanol as mentioned in step 2, freshly prepared clearing solution (preparation described in materials section) containing chloral hydrate is added to the sample. The samples are incubated in the clearing solution for atleast 3 hours prior to mounting the slides for brightfield imaging. Note: Increasing the duration of clearing can enhance the contrast during brightfield imaging to some extent.

### Slide preparation

Using a small paint brush, each cleared leaf is placed on a clean slide with the adaxial surface of the leaf facing upward. The brush can be used to gently tease open any curled leaves without inflicting any further damage. The coverslip is mounted over the sample, taking care not to create any bubble. Multiple leaves (6-8) can be placed under a single coverslip. The clearing solution is used for mounting the samples.

### Brightfield Imaging

The regeneration of vascular strands in the cleared samples can be assessed using brightfield mode in fluorescent or confocal microscope. However, confocal microscope is recommended to acquire high resolution images of the regenerating xylem elements. The settings described here are for Leica TCS SP5 II inverted microscope. Argon laser or DPSS 561 can be used for the imaging at a laser power of 30%, scan speed:200Hz, line average: 2 and pixel format:1024×1024. Newly formed vascular strands display distinct morphology characterised by end-to-end connected xylem elements (Fig1. E,F,J,K,Q).When the regenerating vein re-united the cut ends of the midvein forming a D-loop (Fig1. C, E) or connected either of the cut ends to a lateral vein (Fig1. D,F), the outcomes were scored as successful regeneration. To maintain consistency in the methodology, incisions made in locations other than junction of first lateral vein were not scored while studying age dependency of regeneration. Additionally, only incisions creating a gap less than 400µm between detached parental strands were scored.

### Statistical Analysis

Statistical analysis was performed using R software. The collected data was statistically analysed by Pearson’s χ2 squared test.

## RESULTS AND DISCUSSION

Although the regeneration ability of plants have been widely investigated, leaves are seldom studied for their regeneration and local wound repair ability (Kuchen et al. 2012; Radhakrishnan et al. 2020). We describe a detailed stepwise method of a novel leaf vascular regeneration assay which can be used to study regeneration of the midvein in response to local injury. Our studies have previously shown that a mechanical disconnection of midvein, creating a gap of under 400µm (measured after sample clearing) can be bridged by regenerating vascular strand (Radhakrishnan et al. 2020). While the injured vascular tissue degenerates, the newly synthesized vasculature can either reunite the disconnected strands, or connect the cut end to the nearest lateral vein (Fig. 1B-F). Either way, the reconnection ensures restoration of leaf vascular network, and transport between leaf and rest of the plant body. However, extensive damage generating a gap larger than 400µm cannot be repaired, thereby denying functional restoration of leaf vascular tissue (Radhakrishnan et al. 2020)(Fig.1P). Here, the wound-size dependency of vascular regeneration was recapitulated *in silico* by implementing a computational model based on canalisation hypothesis of vein formation in leaf(Rolland-Lagan and Prusinkiewicz 2005). According to the canalisation hypothesis, positive feedback between auxin flux and PIN1 polarization leads to channelized auxin flow that, in turn, promotes the differentiation of vascular tissue(Tsvi Sachs 1991). Consistent with our previous experimental observations, the computational model demonstrate that the formation of a new vascular strand is indeed dependent on the size of the opening (mimicking wound induced gap) created in a matrix of cells (resembling leaf blade). Equations governing the mathematical model and other relevant details are presented in the supplementary information (Supplementary1, Supplementary Videos). Our result indicates that in addition to higher animal cells and unicellular *Dictyostelium*, wound size sensitivity of the repair process is conserved in plants as well (Pervin et al. 2018).

Having substantiated the wound-size dependency of vascular regeneration, we next investigated whether the regeneration response is dependent on the age of the wounded plant. In many higher animals progressive aging is associated with reduced the regeneration ability (Yun 2015). To probe how age regulates the regeneration response in leaves, we performed the incision in plants of ages ranging from 3dpg to 10dpg. Performing incisions on the miniscule 3dpg leaves were tedious and often damaged the leaves excessively. Upon comparison, 3dpg plants showed lower regeneration efficiency than 5dpg plants (Fig. 1G,1O). Plants belonging to the age group of 4-6dpg displayed highest regeneration efficiency, making it the optimum age to study the vascular regeneration in leaves (Fig.1H-J,1O). Although it is easier to perform incision in older and larger leaves, the regeneration efficiency declined steeply, with 10dpg leaves completely failing to regenerate (Fig.1K-O). It is important to note that even when injury induced gap was lesser than 400µm, vascular regeneration was impeded in these older leaves (Fig.1L-N). Thus our data suggests that vascular regeneration efficiency reduces with increase in the age of the injured plant.

Regeneration studies in plants and animals have demonstrated that the competence to regenerate in response to injuries can vary even within a specific organ (Durgaprasad et al. 2019; Morgan 1902). So we next examined how the position of incision on the growing leaf influenced the vein regeneration efficiency. To analyse this, we made incisions at different positions along the leaf blade, namely, the petiole of the leaf, the basal end (proximal to the plant body axis) of the midvein, the apical end (distal to the plant body axis) of the midvein and on the lateral veins (Fig. 2A, 2C-F). The highest regeneration frequency was recorded at the basal end of the midvein, particularly, between the first and second lateral vein (Fig. 2B, 2D). The petiole also showed similar regeneration efficiency upon incision and often led to formation of multiple strands in response to injury (Fig. 2B, 2C). However, since the leaf is excised at the petiole during sample collection, the incision site and regenerated vascular strand is occasionally damaged leading to loss of valuable samples. Additionally, since incisions performed in the petiole leads to multiple stand formation instead of single stand regeneration, it will be more appropriate to make injuries in leaf blade, as the study involves following a single regenerating strand in real-time to study recognition, communication and reunion of vascular strands. Upon injuring other positions, we observed that the regeneration efficiency drastically declined towards the apical regions of the midvein and in the lateral veins (Fig. 2B, 2E-G).

Collectively, our data demonstrates that the leaf vascular regeneration is sensitive to the size of the wound, the age of the injured leaf and the position of incision on the leaf.

## CONCLUSION

Our study reveals that 4-6dpg leaves respond most efficiently to smaller wounds (400µm or less in size) that are inflicted at the junction of first lateral vein at the proximal end of the leaf blade. While adopting this assay to study regeneration in other plant species, we recommend standardization of the method with respect to the above mentioned criteria.

The assay will be helpful in exploring the mechanisms underlying regeneration of vascular tissue in growing leaves. To begin with, the assay may prove tedious, however with repeated practice; this method can be performed deftly and rapidly in large number of samples. The short duration of the experiment (experimental data can be collected 5 days post injury) and the dispensability of specialized equipment make it amenable to the larger scientific community. Our method, revealing the dependency on leaf age and wound position during vascular regeneration, can be used in combination with other cell biology and molecular biology techniques with little or no standardisation, thereby adding to its utility.

## Supporting information

Video1:Vein formation:No Incision

Video2:Small incision in leaf lamina

Video3:Large Incision in leaf lamina

## ACKNOWLEDGMENT

K.P. acknowledges grants from the Department of Biotechnology (DBT), Government of India [grant BT/PR12394/AGIII/103/891/2014] and Department of Science and Technology, Science and Engineering Research Board (DST-SERB), Government of India [grant EMR/2017/002503/PS] and also acknowledges the Indian Institute of Science Education and Research Thiruvananthapuram (IISER-TVM) for infrastructure and financial support. D.R. and M.M.M acknowledge University Grants Commission (UGC) fellowship. A.K. was supported by Indian Institute of Science Education and Research-Thiruvananthapuram fellowship. A.P.S. and V.V. are recipients of Council of Scientific and Industrial Research (CSIR) fellowships. M.A. acknowledges Department of Biotechnology (DBT), Ministry of Science and Technology, Government of India for granting the DBT-Post Doctoral Fellowship (DBT-RA Program). K.R.M. and R.K.R are funded by DBT. A.S. acknowledges the support of the Science and Engineering Research Board, Government of India through EMR grant No. EMR/2016/007221.We are thankful to Aswathy Syam, Aleesha Jaleel and Kaustuv Ghosh for assistance with preliminary experiments.

## BIBLIOGRAPHY

Durgaprasad, Kavya et al. 2019. “Gradient Expression of Transcription Factor Imposes a Boundary on Organ Regeneration Potential in Plants.” Cell Reports.

Hwang, Hau-Hsuan, Manda Yu, and Erh-Min Lai. 2017. “ Agrobacterium-Mediated Plant Transformation: Biology and Applications.” The Arabidopsis Book.

Ikeuchi, Momoko, Yoichi Ogawa, Akira Iwase, and Keiko Sugimoto. 2016. “Plant Regeneration: Cellular Origins and Molecular Mechanisms.” Development (Cambridge).

Iwase, Akira et al. 2011. “The AP2/ERF Transcription Factor WIND1 Controls Cell Dedifferentiation in Arabidopsis.” Current Biology.

Kareem, Abdul et al. 2015. “PLETHORA Genes Control Regeneration by a Two-Step Mechanism.” Current Biology 25(8): 1017–30. http://dx.doi.org/10.1016/j.cub.2015.02.022.

Kuchen, Erika E. et al. 2012. “Generation of Leaf Shape through Early Patterns of Growth and Tissue Polarity.” Science.

Morgan, T. H. 1902. “Further Experiments on the Regeneration of the Tail of Fishes.” Archiv für Entwicklungsmechanik der Organismen.

Pervin, Mst Shaela et al. 2018. “A Study of Wound Repair in Dictyostelium Cells by Using Novel Laserporation.” Scientific Reports.

Radhakrishnan, Dhanya et al. 2020. “A Coherent Feed-Forward Loop Drives Vascular Regeneration in Damaged Aerial Organs of Plants Growing in a Normal Developmental Context.” Development (Cambridge, England) 147(6): 1–10.

Rolland-Lagan, Anne-Gaëlle, and Przemyslaw Prusinkiewicz. 2005. “Reviewing Models of Auxin Canalization in the Context of Leaf Vein Pattern Formation in Arabidopsis.” The Plant Journal 44(5): 854–65.

Sachs, T., and M. Hassidim. 1996. “Mutual Support and Selection between Branches of Damaged Plants.” In Vegetatio,.

Sachs, Tsvi. 1991. “Cell Polarity and Tissue Patterning in Plants.” Development 113(Supplement 1).

Shanmukhan, Anju Pallipurath et al. 2020. “Regrowing the Damaged or Lost Body Parts.”Current Opinion in Plant Biology.

Yun, Maximina H. 2015. “Changes in Regenerative Capacity through Lifespan.” International Journal of Molecular Sciences.

