## Supplementary Information for "Age, wound size and position of injury – dependent vascular regeneration assay in growing leaves"

### SUPPLEMENTARY1: NUMERICAL SIMULATIONS OF VASCULAR REGENERATION

The *in-Silico* investigations were carried out with the aim of obtaining both quantitative and qualitative understanding of the dependence of vascular regeneration on the size of the wound that was observed in our experiments. We choose to use a well-studied mathematical model that incorporates the feedback between auxin, its flux and active transport (Mitchison, 1981, Rolland-Lagan and Prusinkiewicz 2005) without making any modifications to it. This is to ensure that the focus remains on understanding the experimental observations without deviating into an exploration of the consequences of such modifications, if any. The variables and parameters associated with each cell that appears in the model are schematically shown in Fig.S1.

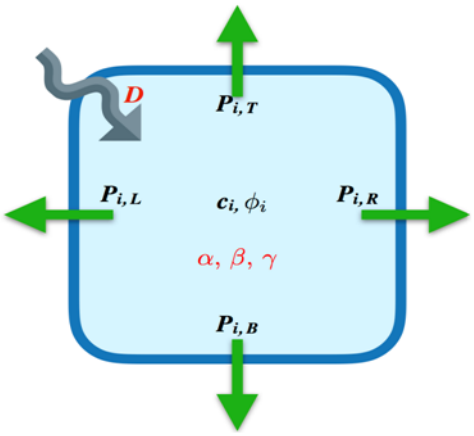


**Figure S1.Schematic of each cell with the variables and parameters appearing in the mathematical model**

Each individual cell is modelled as rectangular or square units and shown in the figure is the $i^{th}$ cell from a large grid. The concentration of auxin in this cell is labelled as $c_I$ while $\phi_I$ denotes the total flux of auxin through the $i^{th}$ cell. Net auxin efflux corresponds to $\phi_I$ having positive values while negative values of $\phi_I$ corresponds to net auxin influx. The total flux is the sum of the fluxes through all four sides of the square cell:

$$\phi_i = \phi_{i,L} + \phi_{i,R} + \phi_{i,B} + \phi_{i,T}, \qquad (1)$$

where the subscripts $L$, $R$, $B$ and $T$ stand for fluxes to/from the left, right, bottom and top respectively. The positive feedback between the concentration and flux of auxin and the efflux carriers (and their polarity) is encoded in the active transport coefficients $P_{i,L}$, $P_{i,R}$, $P_{i,B}$ and $P_{i,T}$ associated with the four walls of the $i^{th}$ cell. In addition to facilitated diffusion, ordinary diffusion of auxin across the four cell walls controlled by the diffusion constant $D$ is also present. The auxin flux across each of the walls is then given by

$$\phi_{i,X} = D(c_i - c_j) + P_{i,X}c_i - P_{j,X'}c_j, \qquad (2)$$

where subscripts $X$ and $X'$ can take on values$L$, $R$, $B$ or $T$. In the equation above, $c_j$ is the concentration of auxin in the adjacent cell on the appropriate side. For instance, if we are considering the flux to/from the left for the $i^{th}$ cell, then $c_j$ is the concentration of auxin in the cell immediately to the left of the one under consideration. In this case the subscript $X'$ in $P_{j,X'}$ takes the value $R$ since the net flux to/from the left for the $i^{th}$ cell also depends on the flux of auxin to/from the right for the cell immediately to the left of the one under consideration.

The active transport coefficients are also variables in the model that change in response to the auxin flux. This positive feedback loop is governed by the equation,

$$\frac{dP_{i,X}}{dt} = \alpha\phi_{i,X}^2\Theta(\phi_{i,X}) + \beta - \gamma P_{i,X} \qquad (3)$$

Where$\alpha$, $\beta$ and $\gamma$ are constants that are tunable parameters in the model. Here $\alpha$ controls the positive nonlinear feedback that auxin the flux $\phi_{i,X}$ has on the transport coefficient. This parameter is relevant only when there is efflux and $\phi_{i,X}$ is positive since the term is multiplied by the Heaviside step function $\Theta(x)$which is zero when $x$ is negative and has unit value when $x$ is positive. The parameter $\beta$ controls the background production of efflux carriers and $\gamma$ is the rate of decay of the same.

The concentration of auxin in each cell is governed by the equation

$$\frac{dc_i}{dt} = \sigma - \sum_{X=L,R,B,T}\phi_{i,X}, \qquad (4)$$

Where$\sigma$ is the intrinsic auxin production rate in each cell. The negative sign before the second term in the equation above appears because we have taken auxin efflux to correspond to positive values of $\phi_{i,X}$.We run our numerical simulations on a 50 by 50 grid of square cells using *Matlab*. In fact, it does not matter whether the cells are square or rectangular since all that matters is that each cell has four neighbours. Initially each of the cells are assigned random small values for both $c_I$ and $\phi_{i,X}$. An auxin sink is present at the bottom boundary of the grid so that there is no flux from below for the bottom-most row of cells. However flux to the bottom $(\phi_{i,B} > 0)$ is allowed for these cells so that they can dump auxin into the sink. The left and right sides of the grid have hard wall boundary conditions so that no flux of auxin to or from the left is allowed for the left-most column of cells and no flux to or from the right is allowed for the right-most column of cells. Similarly for all except two cells in the top row, flux of auxin to or from above is not allowed. At two locations along the top row that divides the 50 cell long array into almost equal thirds, two sources of auxin are introduced. For these two cells influx of auxin from the top is fixed at a constant value.

Numerical integration of the set of equations describing the model proceeds as follows. Starting from the random distribution of the values fluxes and concentration of auxin, the active transport coefficients $P_{i,X}$ associated with each cell is updated according to equation (3). Using the updated values of $P_{i,X}$ the fluxes and concentrations associated with each cell are updated using equations (2) and (4) respectively. A small time step of 0.05 in arbitrary units is used for the numerical propagation of the system of equations. For all the simulations done, we have chosen $\beta = \gamma = 0.0005$, keeping the intrinsic production and decay of auxin efflux carriers relatively low in all cells. There is no intrinsic rate of production of auxin since $\sigma = 0$ in all our numerical runs. The diffusion constant is kept at $D$ = 0.3. In Fig. 2SA the result of integrating the equations through 24000 time steps is shown with the feedback coefficient that regulates the production of efflux carriers in response to increased auxin flux $(\alpha)$ is set to the value 0.05. We see that, as reported earlier as well, this system of equations leads to the formation of a vein like structure through which the flow of auxin is canalized.

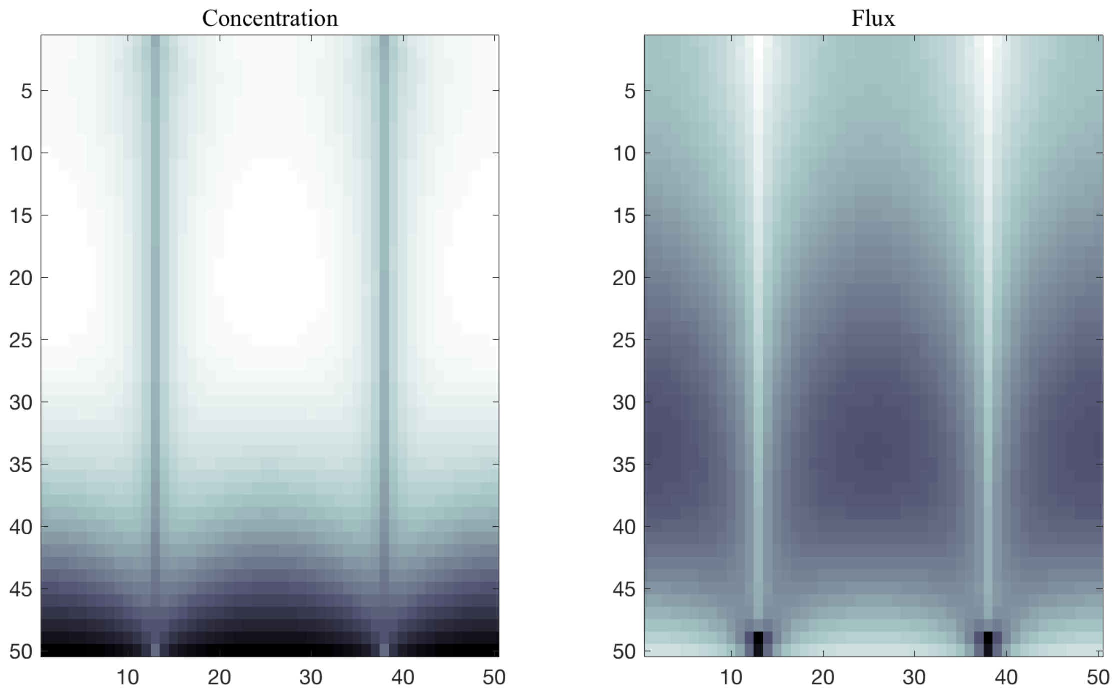


**Figure S2: Formation of veins in a grid of 50 by 50 cells**

In Fig. S2, the panel on the left shows the deviation of the concentration of auxin in each cell from the average. The panel on the right shows the deviation in the total flux of auxin through each cell from the average value. The total flux is computed using equation (1) and the averages as computed across all the cells. In the panel on the left, dark areas correspond to low auxin concentrations relative to the mean and the bright areas correspond to high concentrations relative to the mean. The clearly demarcated vertical lines of cells that start at the locations of the auxin sources can be identified with the veins. Since auxin is being drained quickly in these cells and auxin does not accumulate, we see that the concentration of auxin in the location of the veins is low as expected.

It may be noted that the duration of integration up to 24000 times steps is chosen so that the vein formation has sufficient time to complete connecting the sources at the top to the sink at the bottom. The integration time is long enough to produce a steady pattern. However, the integration cannot be continued much beyond the point after the vein that is dynamically formed connects to the sink. This is because the sink is much stronger than the source and it will drain substantial amount of auxin from the system bringing the concentration down to the level of numerical errors very quickly.

The panel on the right shows the auxin flux. Here dark areas correspond to higher auxin efflux relative to the mean and brighter areas show cells into which there is a net auxin influx. As expected, cells with net auxin influx mark the veins. However the lower tip of the veins that have just made contact with the sink show high auxin efflux. At shorter integration time, the same pattern is seen at the tip of the vein that is forming showing that accumulated auxin at the growing tip leads to an up-regulation of the efflux carriers at the tip.

We next computed what happens when some of the cells in the path of one of the veins (vascular strand on the right in the images) that are formed are removed to simulate the effect of a small incision. A small incision that is 4 cells wide and 2 cells tall was introduced both after the initial formation of the veins as well as before they are formed. Since no qualitative difference is found in the final result, we present the results when the incision is made before the veins develop so that problems associated with very low concentrations of auxin that arise after one of the veins has made contact with the sink are avoided. The result of the numerical simulation is shown in Fig. S3.

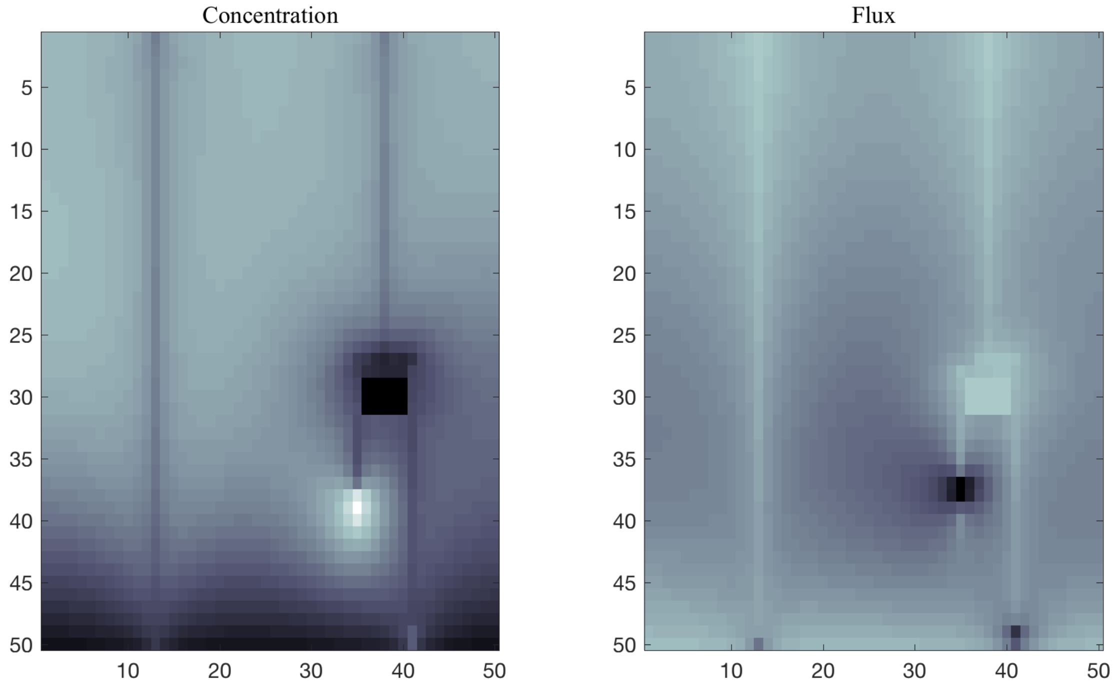


**Figure S3: Formation of veins when there is a small incision**

The boundary conditions set for solving the equations along the edges of the incision are such that there is no flux of auxin into or from the region where the cells have been removed. The incision can be clearly seen in both the panels of Fig. S3. We see that the dynamically formed veins find a path around the small incision and out of the two arms that develop around the incision; one finds its way down to the sink. Once one of the arms reaches the sink, the continued drain of auxin slows down and eventually stops the development of the second

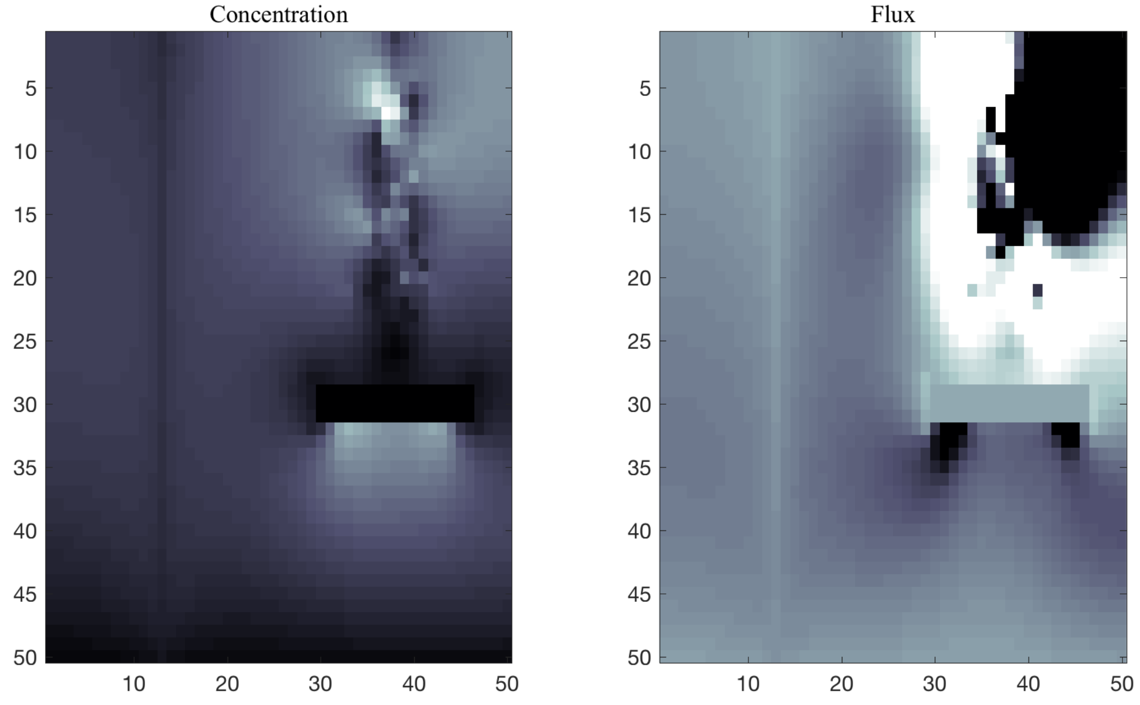


**Figure S4: Development of veins when there is a large incision**

arm. The sink in the realistic case can be another branching vein that has continuous connectivity to the base of the leaf. The feedback control between the auxin flux and the active transport coefficients mimic the relationship between auxin flow and polarized PIN proteins. The numerical model shows that this feedback mechanism can indeed navigate around small incisions and regenerate the vascular tissue (Fig. 1E, 1F).

Fig. S4 shows the development pattern at 24000 time steps when the incision is made much wider at 16 cells wide by 2 cells tall. The feedback mechanism is unable to navigate around the large incision and form a link to the sink. In some isolated cases, a connection to the already formed vein to the left is seen but in general within the time window through which the equations can be numerically integrated without proliferation of numerical errors, the development of the vein is more often than not completely curtailed by the large incision. This is again in support of the observations made in the leaf incision experiments (Fig. 1P).

The pattern of veins produced is relatively robust even if the parameter values are changed provided they are changed proportionately. Increasing or decreasing one of the parameters alone can change the pattern drastically. Thus, here the numerical modelling yields results that reiteratethe experimental observations in response to injury to the vascular tissue in leaves.

### BIBLIOGRAPHY

Mitchison, G.J., 1981. The polar transport of auxin and vein patterns in plants. *Philosophical Transactions of the Royal Society of London. B, Biological Sciences*, *295*(1078), pp.461-471.

Rolland-Lagan, Anne-Gaëlle, and Przemyslaw Prusinkiewicz. 2005. “Reviewing Models of Auxin Canalization in the Context of Leaf Vein Pattern Formation in Arabidopsis.” *The Plant Journal* 44(5): 854–65.

### SUPPLEMENTARY VIDEO FILES

VideoS1: Formation of veins in a grid of 50 by 50 cells with no absent cells (mimicking no incision).

Video S2: Formation of veins in a matrix mimicking small incision.

Video S3: Formation of veins in a matrix mimicking large incision
